# RiboGraph: An interactive visualization system for ribosome profiling data at read length resolution

**DOI:** 10.1101/2024.01.11.575228

**Authors:** Jonathan Chacko, Hakan Ozadam, Can Cenik

## Abstract

**Summary:** Ribosome profiling is a widely-used technique for measuring ribosome occupancy at nucleotide resolution. However, the need to analyze this data at nucleotide resolution introduces unique challenges in data visualization and analyses. In this study, we introduce RiboGraph, a dedicated visualization tool designed to work with .ribo files, a specialized and efficient format for ribosome occupancy data. Unlike existing solutions that rely on large alignment files and time-consuming preprocessing steps, RiboGraph operates on a purpose designed compact file type and eliminates the need for data preprocessing. This efficiency allows for interactive, real-time visualization at ribosome-protected fragment length resolution. By providing an integrated toolset, RiboGraph empowers researchers to conduct comprehensive visual analysis of ribosome occupancy data.

**Availability and Implementation:** Source code, step-by-step installation instructions and links to documentation are available on GitHub: https://github.com/ribosomeprofiling/ribograph. On the same page, we provide test files and a step-by-step tutorial highlighting the key features of RiboGraph.

## Introduction

Ribosome profiling is a widely used method to determine mRNA sequences that are protected from nuclease digestion by ribosomes (Ingolia *et al*., 2009). In combination with RNA expression measurements, ribosome profiling experiments can identify translationally regulated genes across cell types and conditions. This approach has revealed insights into the mechanisms and consequences of translation control in a wide range of organisms and key biological processes (Rao *et al*., 2021; Ozadam *et al*., 2023; McManus *et al*., 2014; Ricci *et al*., 2014; Subtelny *et al*., 2014; Reid and Nicchitta, 2012; Wurth *et al*., 2016; Heyer and Moore, 2016).

Ribosome profiling experiments pose a unique set of challenges for computational analyses due to the information encoded within the variable lengths of ribosome-protected footprints (RPFs). The variation in ribosome lengths is informative for assessing data quality (Cenik *et al*., 2015; Wolin and Walter, 1988; Gerashchenko and Gladyshev, 2016), ribosome conformations (Lareau *et al*., 2014; Wu *et al*., 2019; Guydosh and Green, 2017), and potential collisions (Arpat *et al*., 2020; Meydan and Guydosh, 2020). Conventional methods of storing this information in compressed text or binary alignment files result in substantial storage and computational costs (Cope *et al*., 2022; Ozadam *et al*., 2020). Critically, these conventional formats do not readily support dynamic visualizations at both nucleotide and footprint-length resolutions.

To address these constraints, we introduced a software ecosystem that employs a new hierarchical data structure, termed ‘.ribo,’ to substantially enhance both storage efficiency and data accessibility (Ozadam *et al*., 2020). In the current paper, we extend this framework by describing RiboGraph, a highly interactive, portable and responsive visualization tool that overcomes the limitations of existing alternatives.

While several visualization tools have been described as tailored solutions for ribosome profiling data, many lack interactive capabilities or violate fundamental software engineering practices such as failing to provide source code or installation guidelines. These tools are not considered further. Based on our experience, there are three extant alternative visualization software for ribosome profiling data that provide interactive visualization and conform to usability and best-practices for software development: Shoelaces (Birkeland *et al*., 2018), RiboStreamR (Perkins *et al*., 2019) and riboviz2 (Cope *et al*., 2022).

Shoelaces (Birkeland *et al*., 2018) provides a limited set of visualizations and lacks the capacity to select specific ribosome protected footprint lengths, thereby constraining its utility.

RiboStreamR (Perkins *et al*., 2019) offers a high degree of interactivity but is limited to working with .bam files as input. Consequently, a large amount of preprocessing needs to be completed before working with these files. In our testing, uploading a modest 20-million read .bam file to RiboStreamR’s web server for analyses took approximately 15 minutes per sample, restricting its practical usability.

riboviz2 operates based on an R shiny app and requires a semi-manual installation process involving multiple component packages. In contrast, RiboGraph is readily deployable using a single Docker build command, works with .ribo files, which are orders of magnitude smaller than bam files, and takes only several seconds to start visualization. Concerning user experience, RiboGraph features interactive charts that enable data point hovering as well as chart zooming and panning. riboviz2 provides a less interactive process; the user must drag a slider and press a button to update views and lacks dynamic read-length filtering capabilities. A drawback of riboviz2 is the tight coupling between data preprocessing and visualization, limiting its portability. Specifically, riboviz2 presumes that the user will complete the full analysis pipeline before visualization, thereby limiting its use to a local machine. Conversely, RiboGraph requires no preprocessing of the input files, where input files are small enough to be stored in local compute environments, offers intuitive and interactive visualization and can easily be installed and maintained.

## Implementation and Availability

RiboGraph aims to provide a user-friendly and efficient platform for visualizing ribosome profiling data (Figure 1). We designed this solution to be deployed as either a web service or a local application on the user’s computer. The system operates within Docker containers to ensure consistent performance across different computing environments. For the back end, we employed Django, a popular Python web framework selected for its reliability and flexibility. Instead of using a conventional database, the system integrates with a lightweight SQLite file for lower overhead. On the frontend, Vue.js, a reactive JavaScript framework, is used to dynamically update visualizations based on user selections (Figure 1).

**Figure 1.**
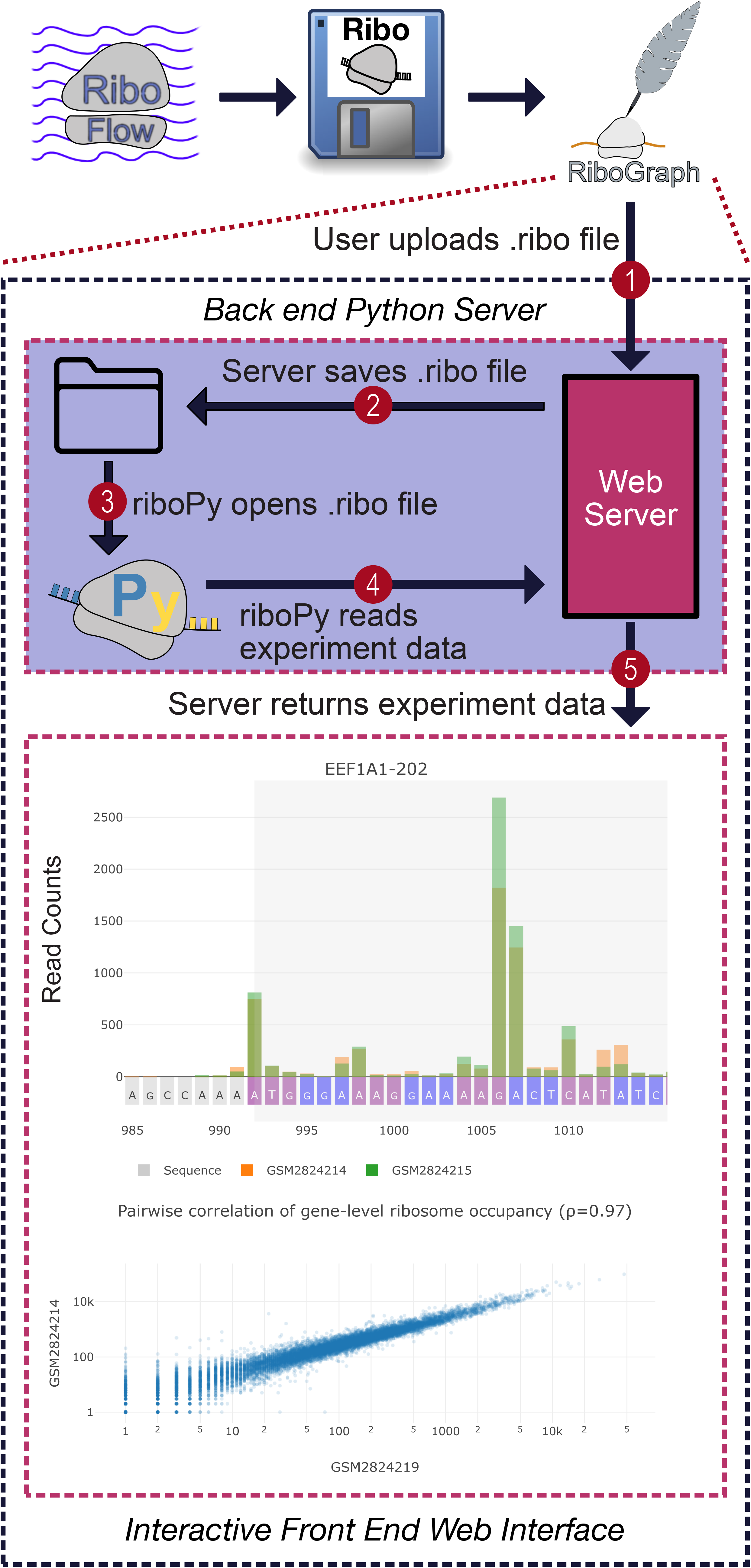
Overview of the RiboGraph visualization software. The ribosome profiling data at nucleotide and footprint length resolution is provided using .ribo files. Experimental data is displayed through an interactive interface to enable commonly used analyses of ribosome profiling at the level of individual experiments or in a comparative manner.

Users can upload .ribo files via a web interface. Upon upload, RiboGraph locally stores these files and creates a corresponding entry in the database, effectively linking each experiment to its data source. When the user accesses a specific experiment’s visualization page, their browser queries the back end server for the necessary data using an HTTP GET request. RiboGraph fetches this file path for the queried experiment from the database and reads the corresponding .ribo file using RiboPy. After performing data cleaning, the system encapsulates the data into a compressed JSON response, which is then sent to the frontend. To expedite subsequent data retrievals, requests are cached on both the frontend and back end, enabling near-instantaneous responses for repeated queries within a short timeframe.

Once the frontend receives the data, it performs client-side filtering and scaling as needed, eliminating repeated server queries and ensuring real-time responsiveness to user actions. Specifically, operations such as footprint length filtering and P-site correction are dynamically executed in the browser using user-selected offsets. The processed data is subsequently visualized through the Chart.js and plotly.js libraries. Utilizing the Vue framework facilitates updating charts in response to user input changes and enhances development modularity through component-based architecture. Collectively, these architectural design choices optimize the distribution of computational load between the frontend and back end, targeting both user experience and system efficiency.

RiboGraph offers comprehensive visualization capabilities to address common tasks in ribosome profiling data analysis (Figure 1). For each experiment, users can explore various facets beginning with the visualization of the distribution of ribosome footprint lengths. Interactive sliders enable dynamic examination of quality control metrics and visualizations across a user-specified range of footprint lengths. The platform supports several common quality control analyses, such as metagene plots that depict ribosome occupancy around start or stop codons aggregated across genes, and ribosome footprint count distributions across different transcript regions like 5′ UTRs, CDSs, or 3′ UTRs. Users have the flexibility to aggregate data or inspect individual transcript coverage, based on desired footprint length ranges. Users can augment individual transcript coverage plots with sequence data by providing a FASTA file. Additionally, users can refine coverage plots through P-site adjustments, either relying on an automated detection algorithm or manually setting offsets.

Importantly, RiboGraph also facilitates cross-experiment comparisons. Users can easily select subsets of experiments for side-by-side analyses. An integrated feature for read-depth normalization permits the direct juxtaposition of experiments, even when they differ in sequencing depth. RiboGraph provides tools to assess reproducibility and data similarity between experiments using adjacency matrix heatmap representing the Spearman correlations between gene-level quantifications. Users can quickly identify outlier genes or patterns between two experiments through interactive scatter plots of the correlation between gene-level transcript footprint counts.

## Conclusion

We introduce RiboGraph, a comprehensive visualization tool designed specifically for ribosome profiling data. RiboGraph not only eliminates the need for bulky files and cumbersome preprocessing steps but also allows for highly interactive, dynamic analyses at ribosome footprint length resolution. The software employs Django for robust back end support and Vue.js for an intuitive front-end experience. Its Docker-based architecture ensures easy installation and consistent performance across platforms. RiboGraph offers a rich set of visualization capabilities including dynamic selection of read-lengths, P-site correction, and direct comparisons across multiple datasets. Moreover, its modularity allows it to serve as a standalone server or integrate into existing software ecosystems. Taken together, RiboGraph will empower researchers as a pivotal tool for efficient analyses of ribosome occupancy data.

## Acknowledgements

The authors acknowledge the Texas Advanced Computing Center (TACC) at The University of Texas at Austin for providing high performance computing and storage resources that have contributed to the research results reported within this paper. URL: http://www.tacc.utexas.edu. All the original text in this paper was written by the authors. A LLM was used to suggest edits for clarity and grammar. We would like to thank Dr. Steve Vokes, Dr. Nimit Jain, Dr. Shilpa Rao, Dr. Vighnesh Ghatpande, Dr. Shilpa Rao, Reiko Tachibana, and Dayea Park for software testing and suggestions for features to include in RiboGraph.

